# Autophagy Decreases Alveolar Epithelial Cell Injury by Suppressing the NF-κB Signaling Pathway and Regulating the Release of Inflammatory Mediators

**DOI:** 10.1101/328039

**Authors:** Tao Fan, Shuo Yang, Zhixin Huang, Wei Wang, Shize Pan, Yao Xu, Boyou Zhang, Zhangfan Mao, Yifan Fang, Xiaobo Guo, Hao Hu, Qing Geng

## Abstract

To research the impact of autophagy on alveolar epithelial cell inflammation and its possible mechanism in early stages of hypoxia, we established a cell hypoxia-reoxygenation model and orthotopic left lung ischemia-reperfusion model. Rat alveolar epithelial cells stably expressing GFP-LC3 were treated with an autophagy inhibitor (3-methyladenine, 3-MA) or autophagy promoter (rapamycin), followed by hypoxia-reoxygenation treatment at 2, 4 and 6h in vitro. In vivo, twenty-four male Sprague-Dawley rats were randomly divided into four groups (model group: no blocking of hilum in the left lung; control group: blocking of hilum in the left lung for 1h with DMSO lavage; 3-MA group: blocking of hilum in the left lung for 1h with 100ml/kg of 3-MA (5μmol/L) solution lavage; rapamycin group: blocking of hilum in the left lung for 1h with 100ml/kg of rapamycin (250nmol/L) solution lavage) to establish an orthotopic left lung ischemia model. This study demonstrated that rapamycin significantly suppressed the NF-κB signaling pathway, restrained the expression of pro-inflammatory factors. A contrary result was confirmed by 3-MA pretreatment. These findings indicate that autophagy reduces ischemia-reperfusion injury by repressing inflammatory signaling pathways in the early stage of hypoxia in vitro and in vivo. This could be a new protective method for lung ischemia-reperfusion injury.

## Introduction

Ischemia-reperfusion (I/R) inflammatory injury, which is characterized by free radical reaction, intracellular calcium overload and leukocyte activation, is a major predisposing factor for lung failure and sudden death in lung transplant operations. However, although intensive investigations of I/R injury in recent decades have promoted the identification of a series of cellular pathologies and improved the operation and survival rate of lung transplantation, many of the mechanisms have not been clarified. Therefore, a better understanding of the pathogenesis of I/R inflammatory injury and the identification of novel therapeutic methods are greatly needed.

I/R injury during lung transplantation involves the induction of genes associated with a number of cellular functions, including apoptosis, inflammation, and oxidative stress(1–3). After pretreatment with I/R, alveolar epithelial cells release inflammatory mediators such as reactive oxygen species (ROS), nitric oxide (NO), tumor necrosis factor α (TNF-α), interleukin-1β (IL-1β) and interleukin-10 (IL-10)(4–9). Nuclear factor kappa B (NF-κB) is a transcription factor that is widely known to be associated with inflammatory responses following ischemia(10, 11). In the early stage of I/R, the activation of IκB kinase beta (IKKβ), the most important kinase upstream of NF-κB, results in the phosphorylation and proteolysis of IκBα, which promote the expression of pro-inflammatory cytokines such as TNF-α and IL-1β(12). The positive feedback cascade in I/R leads to an excessive inflammatory response in the lung, which is the main cause of early complications in patients after lung transplantation. Therefore, blocking the NF-κB signaling pathway is an effective strategy for reducing inflammatory injury during lung I/R(13, 14).

TNF-α, IL-1β, ICAM-1 and MCP-1 are downstream effects of the NF-κB signaling pathway, which are pro-inflammatory cytokines and can be measured to assess NF-κB activity(14–16). IL-10 is an anti-inflammatory cytokine that has a crucial role in preventing inflammatory and immune response(17, 18). In the present study, these inflammatory mediums were detected to verify inflammation activity.

Autophagy is an intracellular self-digesting pathway that delivers cytoplasmic constituents into the lysosome(19). Autophagy controls the turnover of proteins and organelles within cells to help in survival and longevity of cells in metabolic stress(20, 21). Early research indicated that autophagy could be induced by different conditions, including nutrient deprivation/starvation, oxidative stress, hypoxia, and chemotherapeutic drugs(3, 22–24). Autophagy also plays an important role in innate and adaptive immunity and can be regulated by different cytokines, such as TGF-β or IL-6(25–28).

Autophagy, inflammatory cytokines and NF-κB signaling pathways are all involved in lung I/R inflammatory injury, but few researchers have determined its regulatory mechanism. The purpose of this study is to research the impact of autophagy on alveolar epithelial cell inflammatory injury in the early stage of hypoxia in vitro and in vivo and characterize its mechanism. Using an autophagy inhibitor (3-methyladenine, 3-MA) and autophagy promoter (rapamycin) to regulate autophagy levels, we demonstrate that exogenously enhancing autophagy significantly decreases alveolar epithelial cell inflammatory injury by blocking the NF-κB signaling pathway, attenuating pro-inflammatory cytokine expression and increasing anti-inflammatory cytokine expression. These new findings could be a new protective method in lung ischemia-reperfusion inflammatory injury.

## Results

### Effect of 3-MA and rapamycin on GFP-LC3/CCL149 cell viability

The effect of different concentrations of autophagy inhibitor 3-MA and autophagy promoter rapamycin on GFP-LC3/CCL149 cell viability was detected by MTT assay. As shown in **Fig. 1C**, the cell inhibition rates were 14.3, 18.2 and 48.7% for 3-MA at 5, 10 and 15μmol/L, respectively. The cell inhibition rates were 2.3, 2.5 and 2.4% for rapamycin at 150, 200 and 250nmol/L, respectively. Therefore, 5μmol/L 3-MA and 250nmol/L rapamycin were chosen for further experiments.

**Fig. 1.**
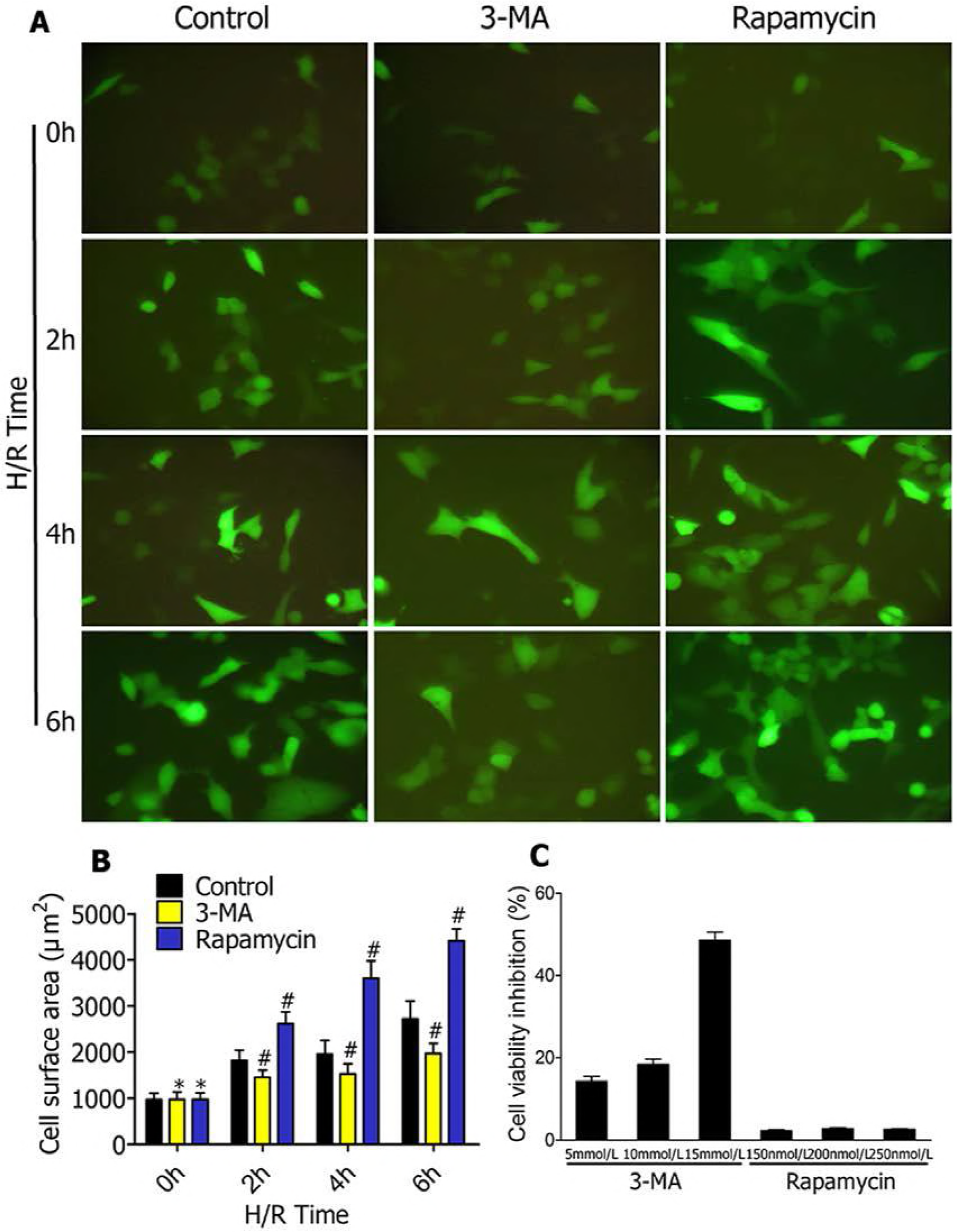
Effect of 3-MA and rapamycin on autophagy formation and GFP-LC3/CCL149 cell viability. **(A)** Representative fluorescence microscopic images of GFP-LC3/CCL149 cells pretreated with 3-MA orrapamycin followed by H/R treatment for 0, 2, 4 and 6h. Scale bars, 40p.m. **(B)** Quantitative results of the green cell surface area of LC3-GFP/CCL149 cells followed by H/R treatment for 0, 2, 4 and 6h in response to DMSO, 3-MA or rapamycin. **(C)** Impact of 3-MA and rapamycin on LC3-GFP/CCL149 cell viability. The cells are treated with different concentrations of 3-MA (5, 10 and 15μmol/L) and rapamycin (150, 200 and 250 nmol/L) for 48h. The control cells are treated with an equal volume of DMSO. MTT assays are used to measure cell viability. The cell inhibition rate (%) is calculated by dividing control values.\**p*≥0.05 compared to control at 0h;#*p*<0.05 compared to control at 2, 4 and 6h.

### Fluorescence microscopy observation

The effect of 3-MA and rapamycin on autophagy formation in GFP-LC3/CCL149 cells was evaluated by observing autophagosomes under fluorescence microscopy following H/R treatment for 2, 4 and 6h **(Fig. 1A)**. Green fluorescence indicated that GFP-LC3/CCL149 cells were successfully constructed. The cellular surface areas of GFP(+) cells were measured by immune staining after pretreating with DMSO, 3-MA (5μmol/L) and rapamycin (250nmol/L) followed by H/R treatment for 0, 2, 4 and 6h **(Fig. 1B)**. Quantitative results of the green cell surface area of GFP (+) cells indicated that 3-MA decreased the expression of autophagy marker protein LC3. In contrast, rapamycin promoted the expression of autophagy marker protein LC3.

### Autophagy is inhibited by 3-MA and strengthened by rapamycin

To research the impact of 3-MA and rapamycin on autophagy, we observed the formation of autophagosomes under transmission electron microscope in GFP-LC3/CCL149 cells after pretreatment with DMSO, 3-MA (5μmol/L) and rapamycin (250nmol/L) followed by H/R treatment for 0, 2, 4 and 6h. As shown in **Fig. 2**, autophagy activity was obviously enhanced in cells pretreated with rapamycin. Autophagosomes are indicated by arrows. We further measured the expression of autophagy-related gene LC3-II/I and Beclin1 in indicated groups. Western blotting results showed the protein levels of LC3-II/I and Beclin1 in GFP-LC3/CCL149 cells **(Fig. 3A)**. The protein levels of LC3-II/I and Beclin1 were quantified and analyzed in the indicated groups**(Fig. 3B, C)**. The protein levels of GFP-LC3 and Beclin1in the 3-MA group were significantly lower than in the control group and those in the rapamycin group were significantly higher than in the control group.

**Fig. 2.**
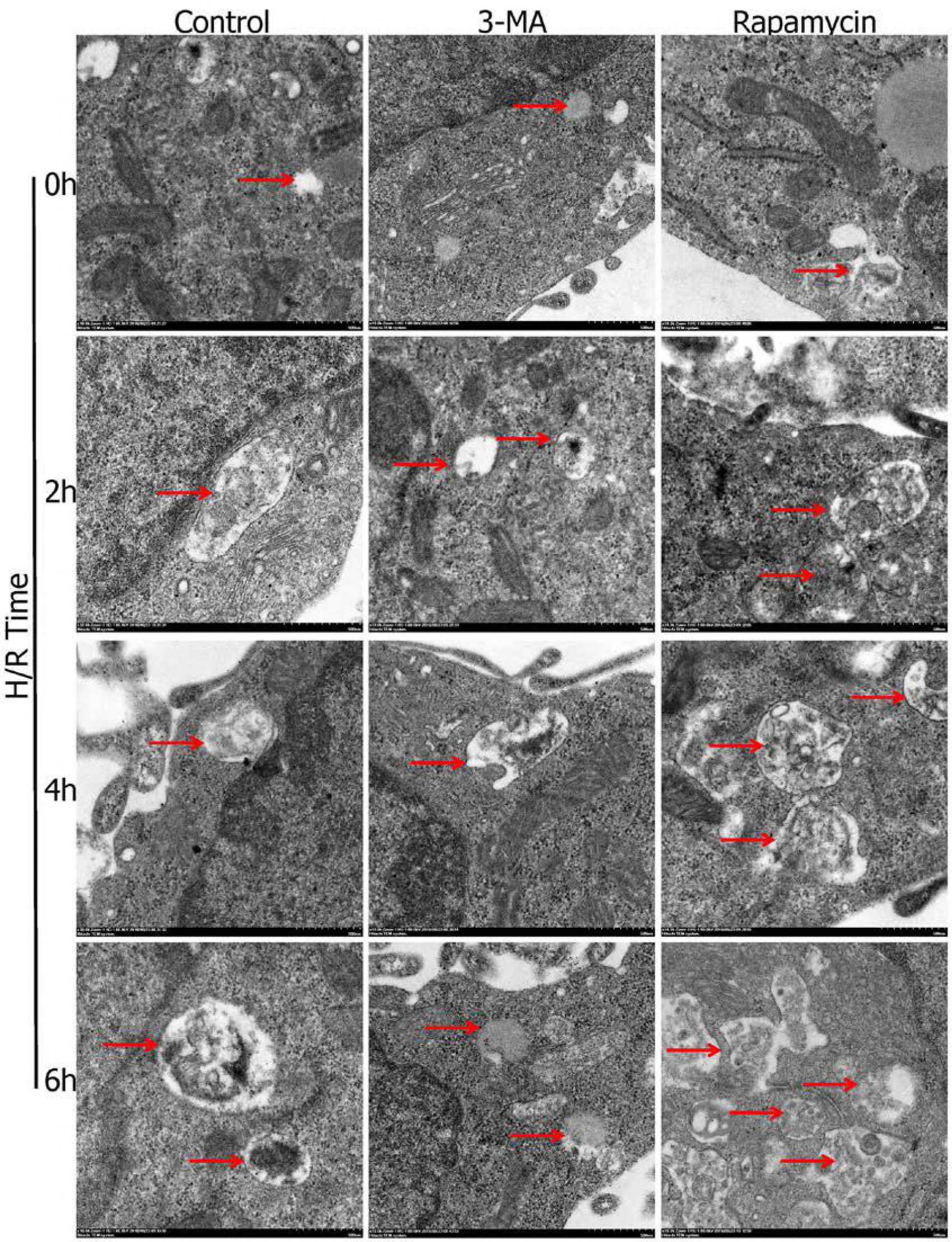
Transmission electron microscope evaluating the effect of 3-MA and rapamycin on autophagosomes in alveolar epithelial cells treated with H/R. The cells are pretreated with 3-MA (5μmol/L) or rapamycin (250nmol/L) followed by H/R treatment for 0, 2, 4 and 6h. Scale bars, 500nm. The cell ultrastructure is observed under a transmission electron microscope. Arrowheads point to intracellular autophagy.

**Fig. 3.**
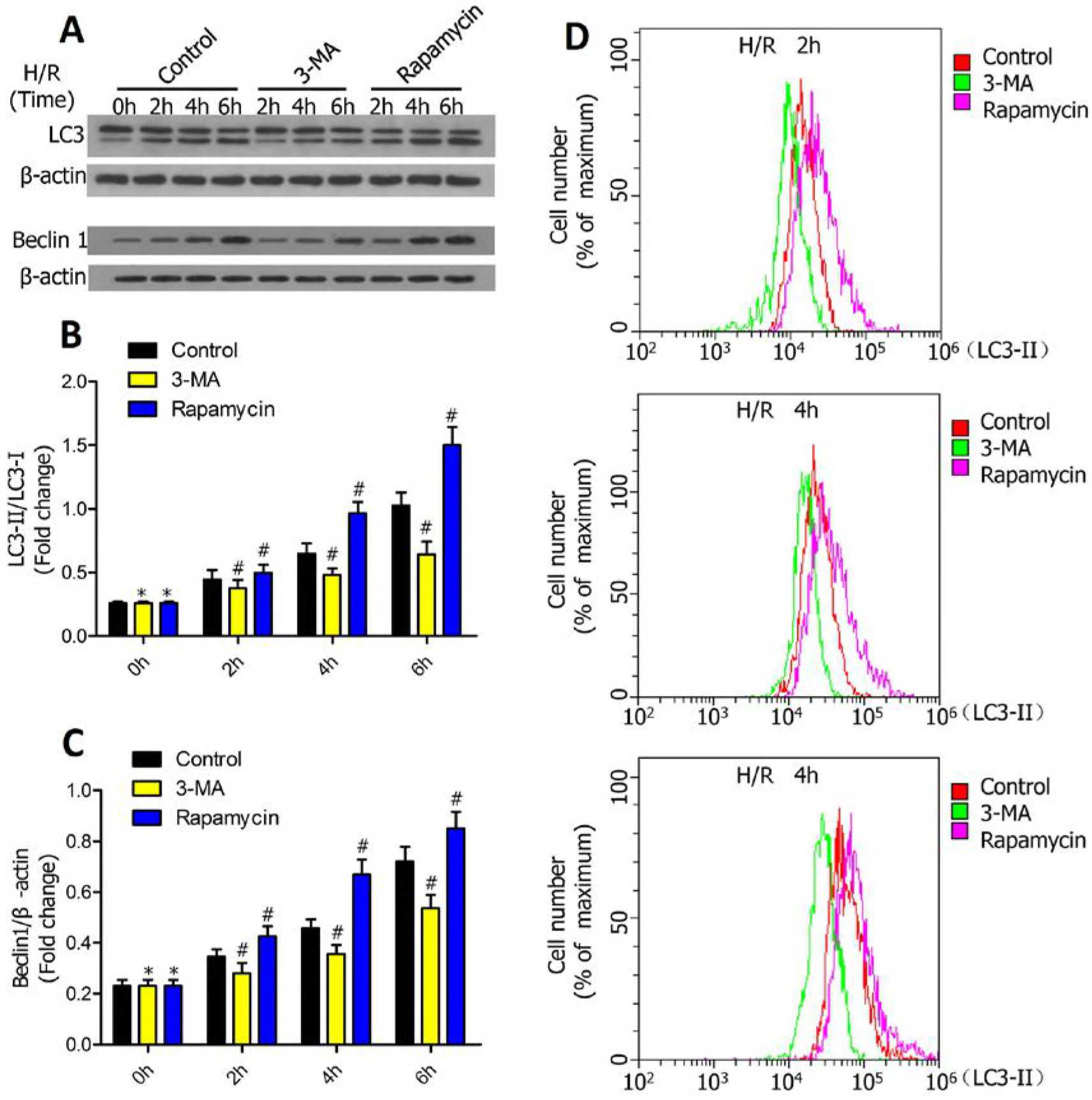
Effect of 3-MA and rapamycin on expression of LC3-II/I and Beclin1 in alveolar epithelial cells treated with H/R. **(A)** Western blots showing the protein of LC3-II/I and Beclin1 in LC3-GFP/CCL149 cells pretreated with DMSO, 3-MA and rapamycin followed by H/R treatment for 0, 2, 4 and 6 h. **(B, C)** The protein levels of GFP-LC3(b) and Beclin1(c) in LC3-GFP/CCL149 cells were quantified and analyzed in the indicated groups. \**p*≥0.05 compared to control at 0h;#*p*<0.05 compared to control at 2, 4 and 6h. **(D)** Flow cytometry was used to measure flux of endogenous LC3 protein. Pretreated with rapamycin exerted a higher level of EGFP-LC3-II-containing autophagosomes.

To further verified the effect of 3-MA and rapamycin on autophagy, we used flow cytometry assay to assess LC3-II in GFP-LC3/CCL149 cells after pretreatment with DMSO, 3-MA (5μmol/L) and rapamycin (250nmol/L) followed by H/R treatment for 0, 2, 4 and 6h. As shown in **Fig. 3D**, the trend of the results was consistent with that of western blot assay and transmission electron microscope assay. The percentage of cells with endogenous LC3 in the rapamycin pretreated group was significantly increased compared to that of the DMSO group and 3-MA group.

### NF-κB was repressed by an autophagy promoter and enhanced by an autophagy inhibitor at an early stage of GFP-LC3/CCL149 cell H/R

To research the impact of autophagy on inflammation at early stages of H/R, we measured the NF-κB signaling pathway by immunohistochemistry in GFP-LC3/CCL149 cells after pretreatment with DMSO, 3-MA (5μmol/L) and rapamycin (250nmol/L) followed by H/R treatment for 0, 2, 4 and 6h **(Fig. 4A)**. Immunohistochemical analysis revealed that NF-κB integrated optical density in 3-MA group was significantly higher than in the control group and in the rapamycin group was significantly lower than in the control group **(Fig. 4C)**. We further measured the protein expression of NF-κB in the indicated groups **(Fig. 4B)**. The protein levels of NF-κB were quantified and analyzed **(Fig. 4D)**. The results suggested that strengthening autophagy suppressed NF-κB protein expression, which indicated that exogenously enhancing autophagy reduced inflammation injury by suppressing the NF-κB signaling pathway in alveolar epithelial cell hypoxia-reoxygenation.

**Fig. 4.**
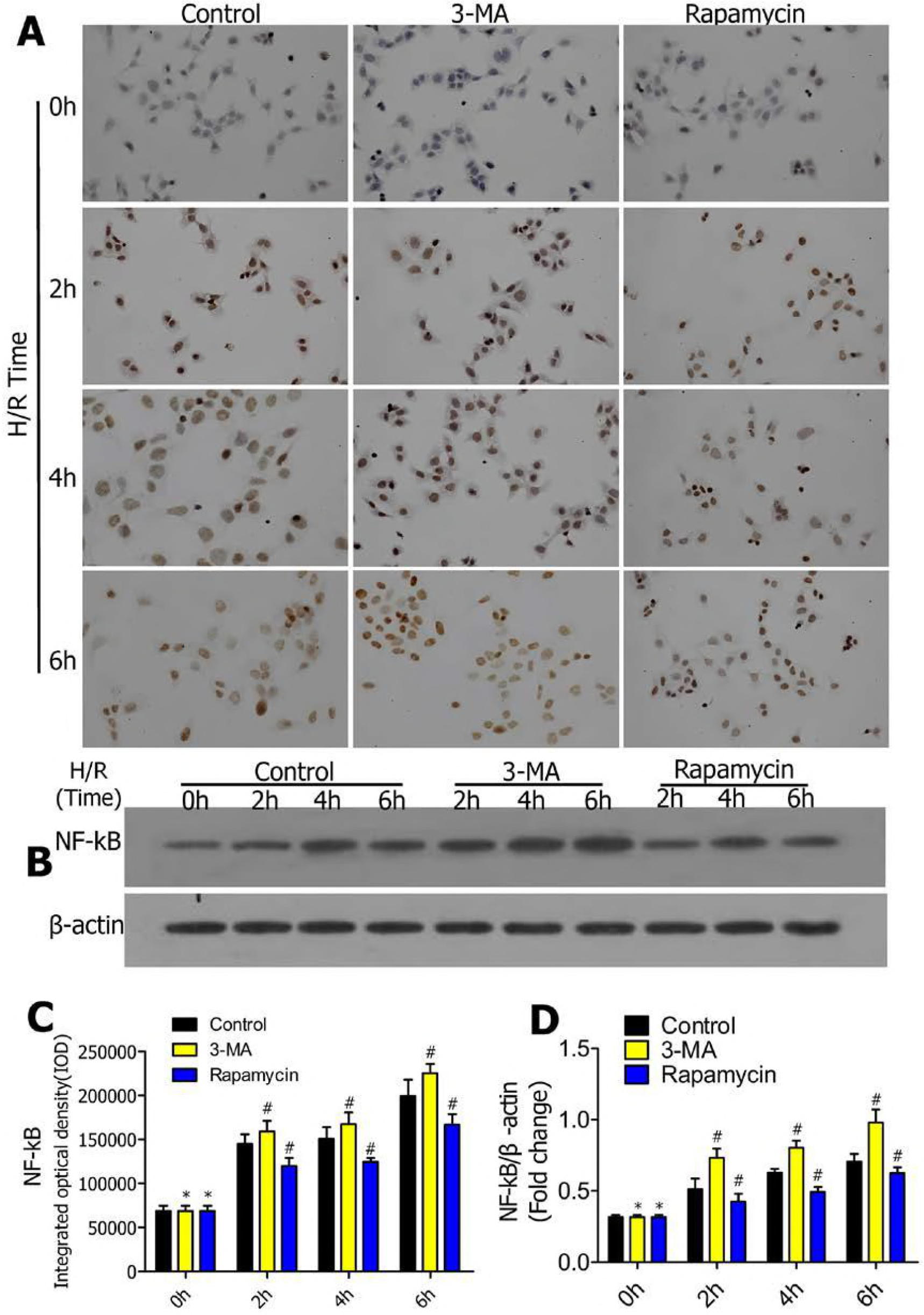
Enhanced autophagy decreases H/R-induced expression of NF-κB in alveolar epithelial cells treated with H/R. **(A)** Representative images of immunohistochemistry with anti-NF-κB antibody in LC3-GFP/CCL149 cells pretreated with DMSO, 3-MA and rapamycin followed by H/R treatment for 0, 2, 4 and 6h(n=5/group; scale bar, 30μm). **(B)** Western blots showing the protein expression of NF-κB in the indicated groups. **(C)** Immunohistochemistry analysis of the protein expressions of NF-κB in LC3-GFP/CCL149 cells in the indicated groups. **(D)** The protein levels of NF-κB in LC3-GFP/CCL149 cells were quantified and analyzed. \**p*≥0.05 compared to control at 0h;#*p*<0.05 compared to control at 2,4 and 6h.

### IκB was enhanced by an autophagy promoter and repressed by an autophagy inhibitor at an early stage of GFP-LC3/CCL149 cell H/R

To further verify the impact of autophagy on the NF-κB signaling pathway at an early stage of H/R, we further detected the IκB expression. We measured IκB by immunohistochemistry in GFP-LC3/CCL149 cells after pretreatment with DMSO, 3-MA (5μmol/L) and rapamycin (250nmol/L) followed by H/R treatment for 0, 2, 4 and 6h **(Fig. 5A)**. IκB integrated optical density in the 3-MA group was significantly lower than in the control group and that in the rapamycin group was significantly higher than in the control group **(Fig. 5C)**. We further measured the protein expression of IκB in the indicated groups **(Fig. 5B)**. The protein levels of IκB were quantified and analyzed **(Fig. 5D)**. The results suggested that strengthening autophagy increased IκB protein expression, which indicated that exogenously enhancing autophagy reduced inflammation injury by increasing IκB expression in alveolar epithelial cell hypoxia-reoxygenation.

**Fig. 5.**
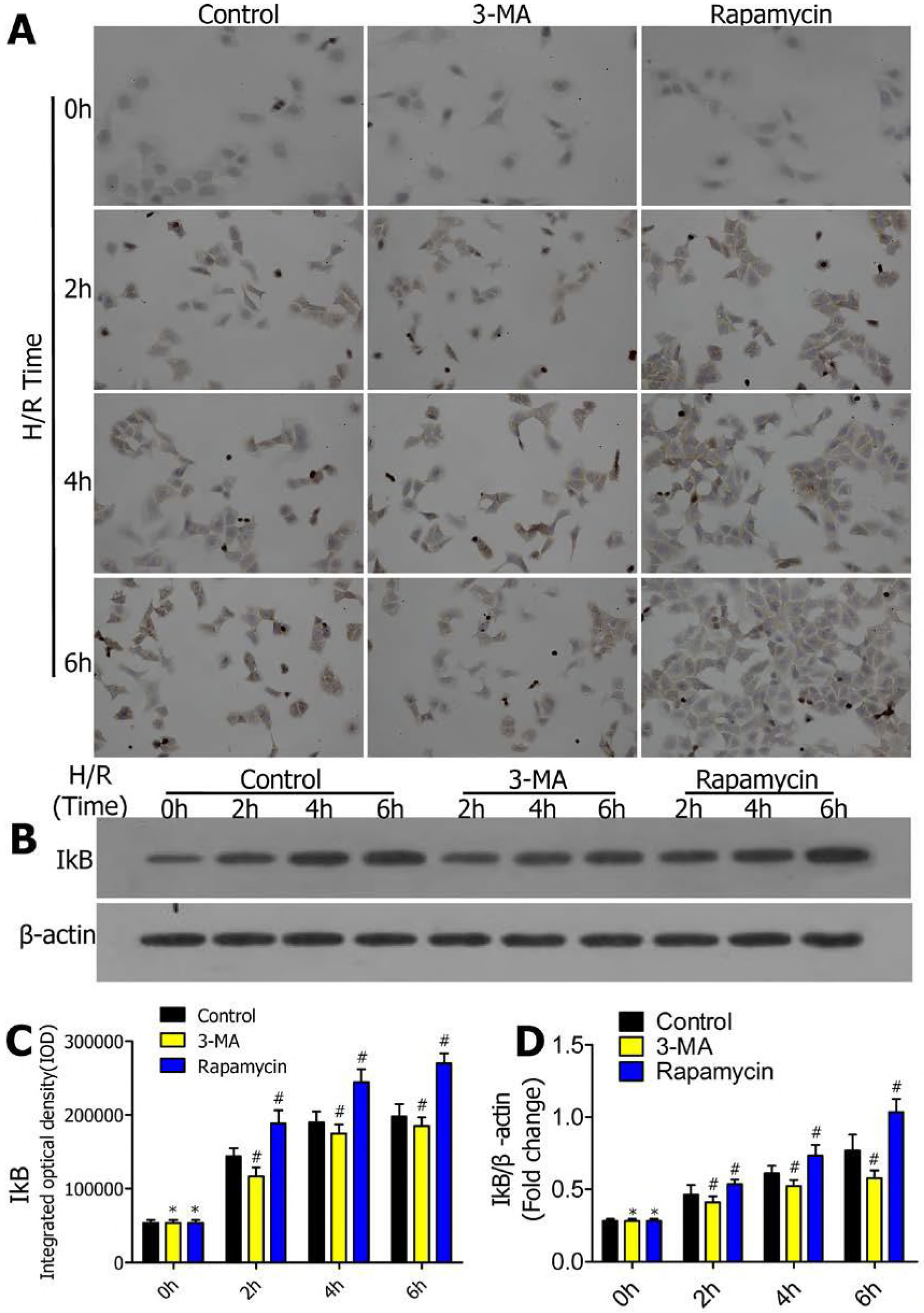
Enhanced autophagy increasesanti-inflammatory factor expression of IκB in alveolar epithelial cells treated with H/R. **(A)** Representative images of immunohistochemistry with anti-IκB antibody in LC3-GFP/CCL149 cells pretreated with DMSO, 3-MA and rapamycin followed by H/R treatment for 0, 2, 4 and 6h(n=5/group; scale bar, 30μm). **(B)** Western blots showing the protein expression of IκB in the indicated groups. **(C)** Immunohistochemistry analysis of the protein expression of IκB in LC3-GFP/CCL149 cells in the indicated groups. **(D)** The protein levels of IκB in LC3-GFP/CCL149 cells were quantified and analyzed. \**p*≥0.05 compared to control at 0h;#*p*<0.05 compared to control at 2,4 and 6h.

### Effect of autophagy on inflammatory factors at an early stage of GFP-LC3/CCL149 cell H/R

In addition to the NF-κB signaling pathway, we also examined the influence of autophagy on downstream effects of the NF-κB signaling pathway in GFP-LC3/CCL149 cells after pretreatment with DMSO, 3-MA (5μmol/L) and rapamycin (250nmol/L) followed by H/R treatment for 0, 2, 4 and 6h. After determining the concentration of each cellular factor using a spectrophotometer **(Fig. 6A, C, E, G, I)**, we measured pro-inflammatory factors TNF-α and IL-1β, MCP-1, ICAM-1 and anti-inflammatory cytokine IL-10 by ELISA kits **(Fig. 6B, D, F, H, J)**,. The result indicated that enhancing autophagy restrained pro-inflammatory factor expression and increased anti-inflammatory cytokine expression.

**Fig. 6.**
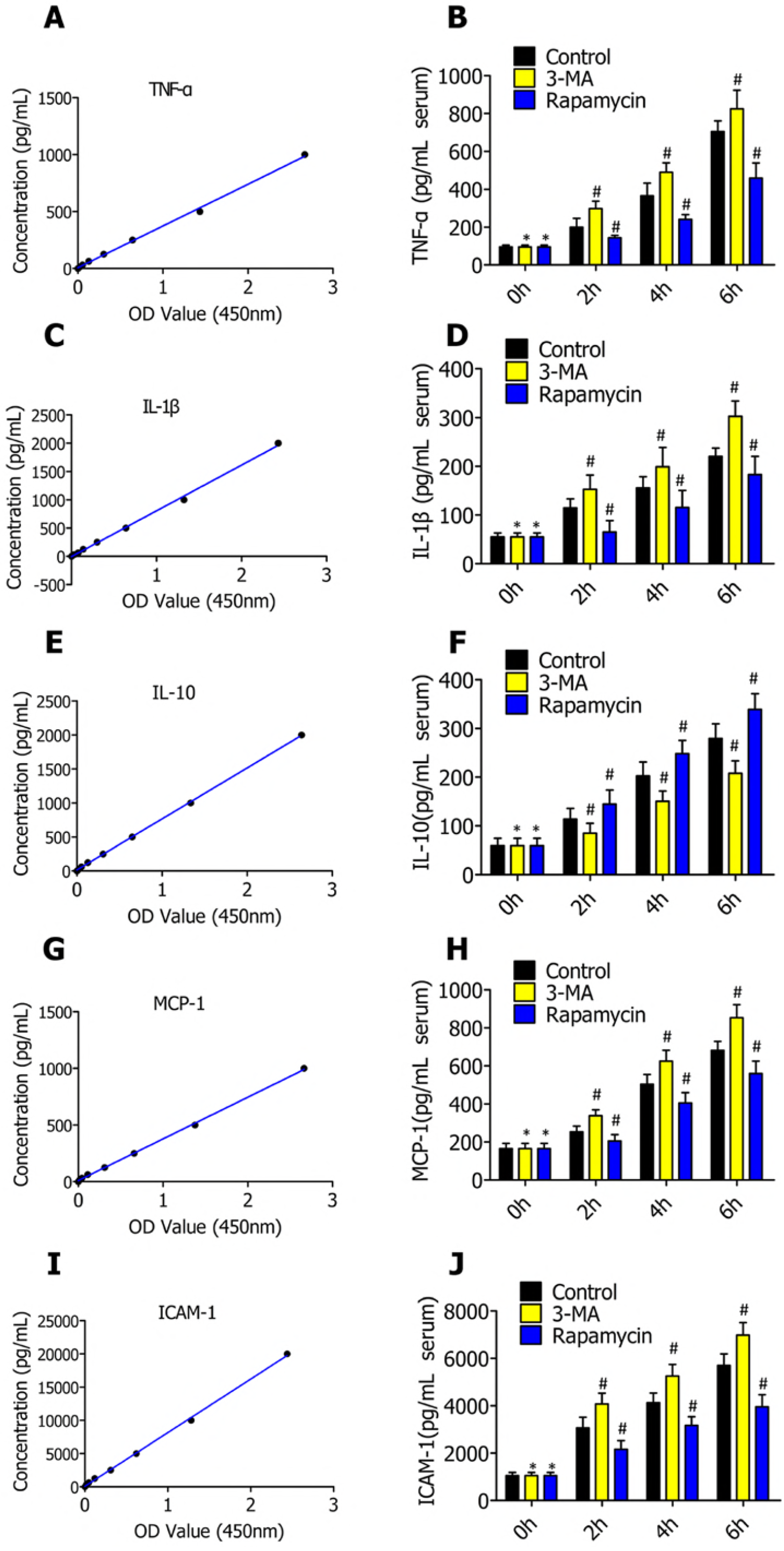
Enhanced autophagy suppresses H/R-induced pro-inflammatory cytokine expression of TNF-α, IL-1β, MCP-1 and ICAM-1and increases anti-inflammatory factor expression of IL-10 in alveolar epithelial cells treated with H/R. **(A, C, E, G, I)** Determining the concentration of TNF-α, IL-1β, IL-10, MCP-1 and ICAM-1 protein concentration in each sample. **(B, D, F, H, J)** ELISA measurement of serum TNF-α, IL-1β, IL-10, MCP-1 and ICAM-1 levels in LC3-GFP/CCL149 cells pretreated with DMSO, 3-MA and rapamycin followed by H/R treatment for 0, 2, 4 and 6h(n=5). The results are analyzed in the indicated groups. \**p*≥0.05 compared to control at 0h;#*p*<0.05 compared to control at 2, 4 and 6h.

### Autophagy activity was inhibited by 3-MA and strengthened by rapamycinin alveolar epithelial cells in rat lung I/R

To further illuminate whether autophagy activity is regulated by 3-MA and rapamycin in alveolar epithelial cells in rat lung I/R, we lavaged rat lungs with DMSO, 3-MA or rapamycin during lung ischemia for 1h and then reperfusion for 2h. The model group was not treated with ischemia. Subsequently, we observed the formation of autophagosomes under a transmission electron microscope **(Fig. 7A)** and detected the expression of autophagy-related gene GFP-LC3 and Beclin1 by Western blotting **(Fig. 7B)**. The protein levels of GFP-LC3 and Beclin1 were quantified and analyzed in the indicated groups **(Fig. 7C, D)**. The protein levels of GFP-LC3 and Beclin1in the 3-MA group were significantly lower than in the control group and those in the rapamycin group were significantly higher than in the control group.

**Fig. 7.**
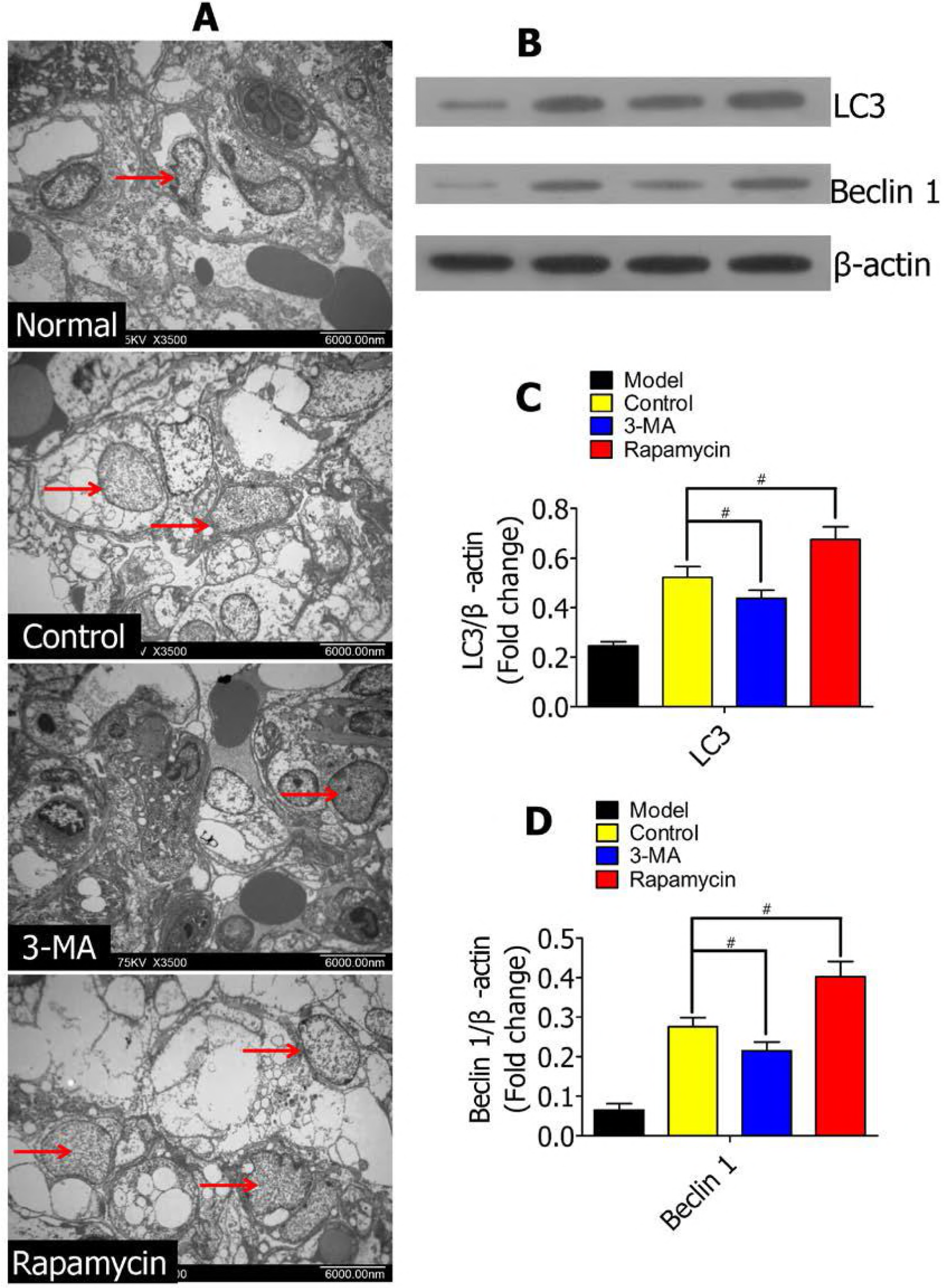
Effect of 3-MA and rapamycin on autophagosome and expression of LC3 and Beclin1in rat lungs treated with I/R. **(A)** Transmission electron microscope image showing autophagosomes in lung tissues from Lewis rats after the rats were lavaged with DMSO, 3-MA and rapamycin in lung ischemia for 2h and reperfusion for 2h. The model group was not treated with ischemia. Arrowheads point to autophagosomes. **(B)** Western blots showing the protein levels of LC3 and Beclin 1 in lung tissues from Lewis rats after the rats were lavaged with DMSO, 3-MA and rapamycin in lung ischemia for 2h and reperfusion for 2h. The model group was not treated with ischemia. **(C, D)** The protein levels of LC3 and Beclin 1 in Lewis rats were quantified and analyzed in the indicated groups.#*p*<0.05 compared to control.

### The NF-κB signaling pathway was restrained by autophagy in alveolar epithelial cell in rat lung I/R

To gain insight into the effect of autophagy on the NF-κB signaling pathway in rat lung I/R, we lavaged rat lungs with autophagy promotor (rapamycin) or inhibitor (3-MA) during lung ischemia for 1h and then reperfused for 2h. The model group was not treated with ischemia. NF-κB and IκB were measured by immunohistochemistry **(Fig. 8A)**. Immunohistochemical analysis revealed that strengthening autophagy suppressed NF-κB protein expression and increased IκB protein expression **(Fig. 8B)**. Western blotting results showed the protein levels of NF-κB and IκB in lung tissues from Lewis rats after pretreatment with I/R **(Fig. 8C)**. The protein levels of NF-κB and IκB were quantified and analyzed in the indicated groups **(Fig. 8D, E)**. These results further revealed that exogenously enhancing autophagy reduced inflammation injury by suppressing the NF-κB signaling pathway in alveolar epithelial cells in rat lung I/R.

**Fig. 8.**
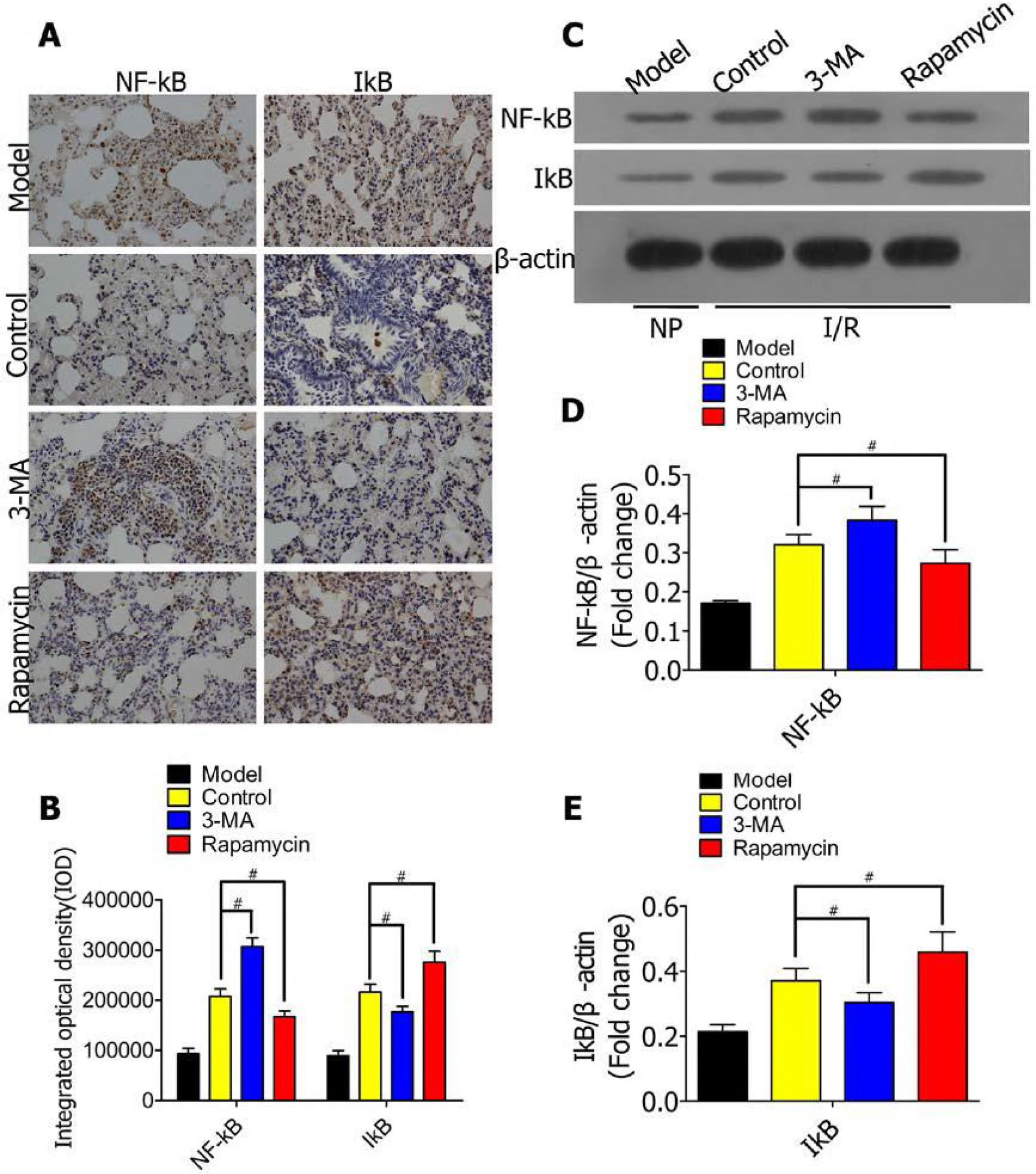
Enhanced autophagy blocks the NF-κB signaling pathway by inhibiting NF-κB expression and increasing IκB expression in rat lungs treated with I/R. **(A)** Representative images of immunohistochemical staining of a normal rat lung section (model group) and lung sections lavaged with DMSO (control group), 3-MA (3-MA group) and rapamycin (rapamycin) with an antibody against NF-κB or IκB; scale bar, 30μm. **(B)** Immunohistochemistry analysis of the protein expression of NF-κB and IκB in rat lungs in the indicated groups. **(C)** Western blots showing the protein levels of NF-κB and IκB in lung tissues from Lewis rats after the rats were lavaged with DMSO, 3-MA and rapamycin in lung ischemia for 2h and reperfusion for 2h. The model group was not treated with ischemia. **(D, E)** The protein levels of NF-κB and IκB in Lewis rats were quantified and analyzed in the indicated groups.#*p*<0.05 compared to control.

### Effect of autophagy on inflammatory factors in alveolar epithelial cells in rat lung I/R

To obtain further knowledge regarding the mechanisms of autophagy reducing alveolar epithelial cell inflammation injury in oxygen deficiency, we detected inflammatory factors in rat lung I/R, which were lavaged with DMSO, 3-MA or rapamycin during lung ischemia for 1h and then reperfusion for 2h. The pro-inflammatory factors TNF-α and IL-1β, MCP-1 and ICAM-1in the 3-MA group were significantly higher than in the control group, and those in the rapamycin group were significantly lower than in the control group **(Fig. 9A, C, D)**. In contrast, anti-inflammatory factor IL-10 showed an increasing trend with enhanced autophagy levels **(Fig. 9B)**. The result indicated that enhancing autophagy can lessen inflammatory injury by restraining the expression of pro-inflammatory factors and increasing anti-inflammatory cytokine expression.

**Fig. 9.**
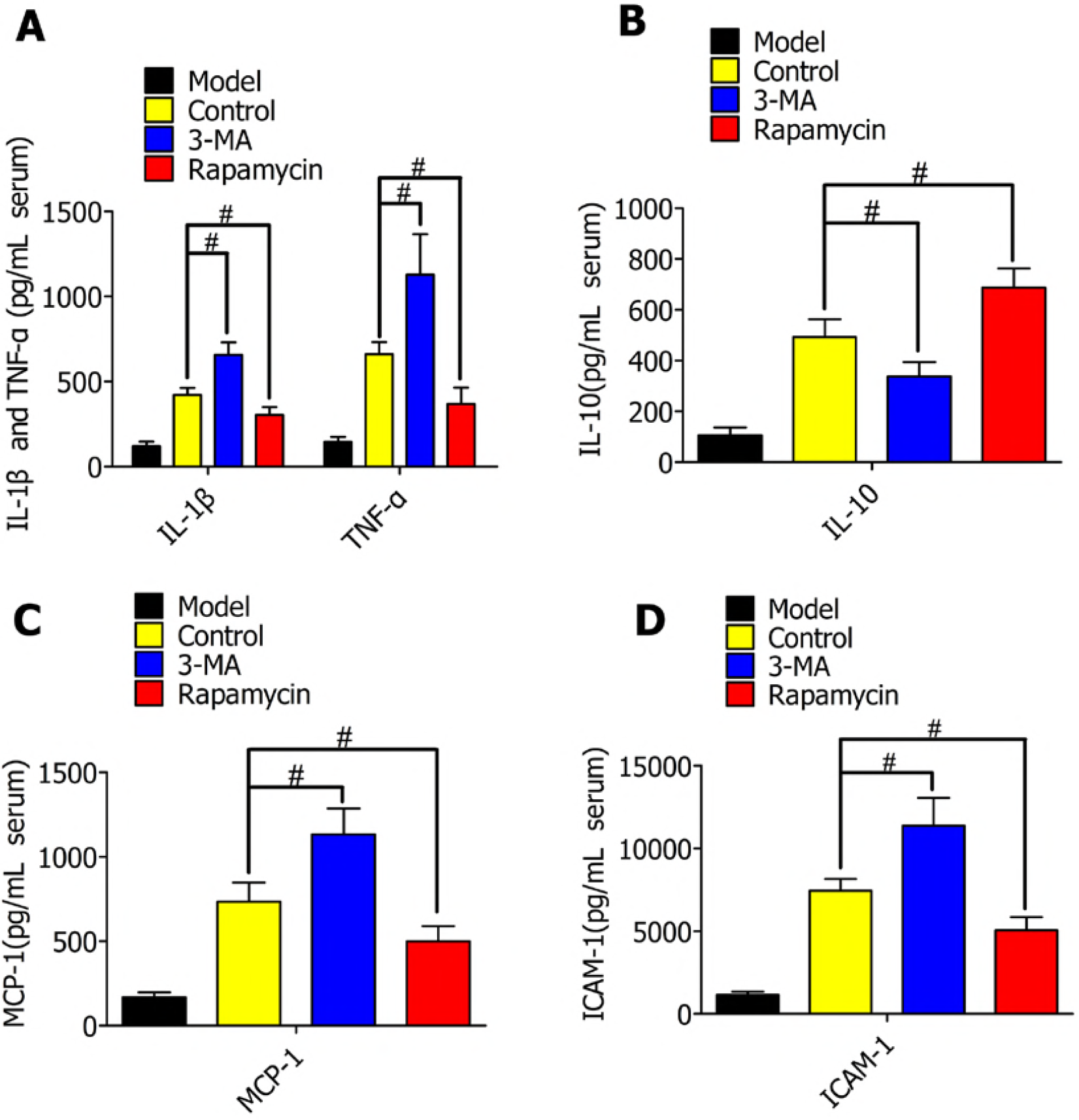
Enhanced autophagy suppresses H/R-induced pro-inflammatory cytokine expression of TNF-α, IL-1β, MCP-1 and ICAM-1and increases anti-inflammatory factor expression of IL-10 in rat lungs treated with I/R. **(A)** ELISA measurement of serum IL-1β and TNF-α levels in rat lungs of control, 3-MA and rapamycin groups (n=5). The results are analyzed in the indicated groups. **(B)** ELISA measurement of serum IL-10 level in rat lungs of control, 3-MA and rapamycin groups (n=5). The results are analyzed in the indicated groups. **(C)** ELISA measurement of serum MCP-1 levels in rat lungs of control, 3-MA and rapamycin groups (n=5). The results are analyzed in the indicated groups. ELISA measurement of serum ICAM-1 levels in rat lungs of control, 3-MA and rapamycin groups (n=5). The results are analyzed in the indicated groups. #*p*<0.05 compared to control.

## Discussion

Lung ischemia reperfusion (I/R) is a frequent event in clinic processes, inducing distant organ dysfunction, especially lung transplantation or acute pulmonary tissue injury. The release of pro-inflammatory cytokines during I/R is one of the most important factors that lead to lung failure(29). Our team has proved that exogenously enhancing autophagy decreased alveolar macrophages apoptosis by attenuating endoplasmic reticulum stress and oxidative stress in hypoxia-reoxygenation or ischemia-reperfusion injury(30). In this study, we established a cell model of alveolar epithelial cell hypoxia-reoxygenation (H/R) and a rat model of focal lung I/R. For the first time, the present study reveals that rapamycin decreases alveolar epithelial cell inflammatory injury by blocking the NF-κB signaling pathway, attenuating the expression of pro-inflammatory cytokines TNF-α, IL-1β, ICAM-1 and MCP-1 and increasing the expression of anti-inflammatory cytokine IL-10. Based on various in vivo and in vitro tissue ischemia and cell hypoxia models, we clearly identified that exogenously enhancing autophagy as a positive regulator of alveolar epithelial cells and lungs responding to oxygen deficiency via blockade of the NF-κB signaling pathway attenuate pro-inflammatory cytokine expression and increase anti-inflammatory cytokine expression.

It has been confirmed that inflammation is an important component of lung I/R. Previous studies demonstrated that inflammatory cytokine infiltration into the lungs during I/R injury participates in the pathogenesis of acute lung failure, especially in patients after lung transplantation. Previous studies have demonstrated that the NF-κB pathway involved in lung disease is induced by I/R(13, 15, 16). NF-κB is combined with the inhibitory unit inhibitory κB (IκB) and is located in the cytoplasm. When IκB is phosphorylated by its kinase IκB kinase (IKK), NF-κB could trigger multiple downstream effects including activation of pro-inflammatory cytokines (TNF-α and IL-1β), ICAM-1 and MCP-1 accumulation, and the infiltration of immune cells in ischemic tissues(31–35). It is also reported that IL-10 can block transepithelial migration of neutrophils(36, 37), which is tightly related to inflammatory and autoimmune pathologies(17). Therefore, inhibiting the inflammatory response is an effective therapeutic method to improve lung I/R injury.

To the best of our knowledge, the present study is the first to show that exogenously enhancing autophagy markedly stimulates the expression of anti-inflammatory cytokine IL-10 at an early stage of hypoxia in vitro and in vivo. Furthermore, the NF-κB signaling pathway and its downstream effects on expression were significantly inhibited by rapamycin under conditions of oxygen deficit. The current results suggested that an autophagy promoter could be a new protective method in lung inflammatory injury induced by ischemia-reperfusion.

## Materials and Methods

### Cell culture

For the in vitro studies, the alveolar epithelial cell line CCL149 (ATCC, Manassas, VA, USA, #CCL149) was chosen as the cell model. The cells were maintained in F-12K medium (ATCC, Manassas, VA, USA)supplemented with 20% fetal bovine serum (FBS, Invitrogen, Carlsbad, CA, USA)at 37°C in a humidified 5%CO_2_ atmosphere. Additionally, 10% heat-inactivated fetal calf serum was contained in the medium. When the cells reached 80% confluence, they were digested with 0.25% trypsin.

### Constructing a stable GFP-LC3/CCL149 cell line

Briefly, the GFP-LC3 plasmid (Addgene, Cambridge, MA, USA) was transfected into CCL149 cells by applying Lipofectamine 2000 reagent (Invitrogen, Carlsbad, CA, USA). The experiment was conducted in accordance with the instructions. Twenty-four hours later, the cells were transferred to culture in F-12K medium containing 300μg/ml of G418 (Invitrogen, Carlsbad, CA, USA). After 2 weeks of expansion, the CCL149 cells were observed under a fluorescence microscope (Olympus, Japan), and the strong green fluorescent colonies were selected as stable GFP-LC3/CCL149 cells and cultured in medium containing 100μg/ml of G418 and 10% FBS for further experiments in the study.

### MTT assay

The general viability of the cells was measured using an MTT assay(38). The percentage of cell viability inhibition was calculated as: cell viability = [OD (treated)−OD (control)]/OD (control) × 100.

### Animal models and procedures

All the animal experimental protocols were approved by the Animal Care and Use Committee of Renmin Hospital of Wuhan University and were conducted in accordance with the National Institutes of Health (NIH) Guide for the Care and Use of Laboratory Animals

Male Sprague-Dawley (SD) rats (8weeks old, 250-300g) were fed a standard diet and maintained in a controlled environment of the animal center. In brief, rats were anesthetized by an intraperitoneal injection of 10% chloral hydrate (300mg/kg body weight) and placed in a supine position. The animals were then intubated for artificial ventilation with oxygen using a small animal breathing machine (5 ml tidal volume, frequency of 70 per min) and electrocardiograph monitoring. Thoracotomy was performed at the anterior lateral side of the left fourth intercostal. The muscular layer and pleura were gently dissected to expose the heart and lung. Then, the hilum of the left lung was dissociated, and an artery clamp was used to pass through the hilum of the lung from the upper right to the lower left. The whole clamped left hilum was clearly exposed by slightly stirring up the clamp. Twenty-four SD rats were randomly divided into four groups (5 rats/group) as follows: (1) model group: no blocking of hilum in the left lung; control group: blocking of hilum in the left lung for 1h with DMSO lavage and then reperfusion for 2h; 3-MA group: blocking of hilum in the left lung for 1h with 100ml/kg of 3-MA (5μmol/L) solution lavage and then reperfusion for 2h; (4) rapamycin group: blocking of hilum in the left lung for 1h with 100ml/kg of rapamycin (250nmol/L) solution lavage and then reperfusion for 2h. Rats in the four groups were sacrificed after the experiment. The left lung tissue of the rats was dissected for further analysis.

### Immunofluorescence analysis

The cell surface area of GFP-LC3/CCL149 cells was assessed by immunofluorescent staining. Briefly, after the hypoxia-reoxygenation (H/R) for 0, 2, 4 or 6 h, the cells were subsequently fixed with 4% paraformaldehyde (Sigma, USA,#158127), permeabilized with 0.1%Triton X-100/BS for 45 min and then stained with β-actin (1:100 dilution),followed by a fluorescent secondary antibody. The surface areas were measured using Image-Pro Plus6.0 software. Images were captured using a special fluorescence microscope (Olympus, Japan).

### Transmission electron microscope

In vitro, cells were fixed with 2.5% glutaraldehyde at 4°C overnight after the H/R treatment for 2, 4 or 6h and then fixed with 1% osmic acid. After being dehydrated with a graded series of ethanol (50, 70, 80, 95, and 100%; each for 15 min) and acetone (twice; each for 15 min), the cells were embedded in epoxide resin.

Ultra-thin sections were generated using an ultra-microtome (LKB-V, Bromma, Sweden) followed by staining with uranyl acetate and lead citrate. Then, sections were observed and photographed under a transmission electron microscope (H-600, Hitachi, Tokyo, Japan).

In vivo, paraffin-embedded lungs were cut transversely into 0.1μm sections. Then, the sections were observed and photographed under a transmission electron microscope (H-600, Hitachi, Tokyo, Japan).

### Flow cytometry

GFP-LC3/CCL149 cells were harvested with DMSO, 3-MA or rapamycin, and washed with PBS containing 0.05% saponin. For intracellular staining of endogenous LC3, CCL149 cells were harvested with trypsin, rinsed with culture medium and PBS, and rinsed with PBS containing 0.05% saponin. Cells were then incubated with mouse anti-LC3 primary antibody (Abcam, Ab290) for 20 minutes, rinsed with PBS, incubated with goat antimouse secondary antibody conjugated to R-Phycoerythrin (BosterBiotech, BA1060) for 20 minutes, and rinsed twice with PBS. More than 30,000 events were captured for every analysis. Fluorescence activated cell sorter data were collected using a fluorescence activated cell sorter Calibur flow cytometer (Becton Dickinson) with Cell Quest Pro software. This method was previously published in ref(39).

### Immunohistochemical analysis

For immunohistochemistry in vitro, GFP-LC3/CCL149 cells growing on glass cover slips were fixed for 15min with 4% paraformaldehyde. After being incubated with 0.5% Triton X-100/PBS solution for 30 min and washed with PBS three times, the GFP-LC3/CCL149 cells were blocked with 3% hydrogen peroxide for 15 min and subsequently incubated overnight at 4 °C with the primary antibodies. Binding was visualized with the appropriate peroxidase-conjugated secondary antibodies (AR1022, ZSGB-BIO) for 20-30 min at 37 °C.

For immunohistochemistry in vivo, paraffin-embedded lungs were cut transversely into 5μm sections. Following a 5min high-pressure antigen retrieval process in 0.1mol/L citrate buffer with a pH of 6.0, the lung sections were blocked with 3% hydrogen peroxide for 15 min and subsequently incubated overnight at 4 °C with the primary antibodies. Binding was visualized with the appropriate peroxidase-conjugated secondary antibodies (AR1022, ZSGB-BIO) for 20-30min at 37 °C.

### Western blotting analysis

Total proteins were extracted from GFP-LC3/CCL149 cells and rat lung tissues in lysis buffer. The protein concentrations were determined using a Pierce Bicinchoninic Acid Protein Assay kit (Biyuntian, Shanghai, China,#P0010). Fifty micrograms of protein was subjected to SDS-polyacrylamide gel electrophoresis (12%PAGE; Amresco) and transferred to a polyvinylidene fluoride membrane (Millipore) followed by incubation overnight at 4 °C with the following primary antibodies: LC3 (Abcam, Cambridge, United Kingdom,#ab62341; 1:200 dilution), Beclin-1 (Santa Cruz, CA, USA, #sc-11427; 1:200 dilution), NF-κB (Bioworld, CA, USA, #BS1257; 1:600 dilution), and IκB-α (Bioworld, CA, USA, #BS3601; 1:600 dilution). After incubation with peroxidase-conjugated secondary antibodies (BA1060, at 1:50,000 dilution), the bands were visualized using Bio-Rad Chemi DocTM XRS+ (Bio-Rad). Protein expression levels were normalized to the corresponding β-actin levels.

### ELISA measurements of cytokines in rat lung tissues and GFP-LC3/CCL149 cells

Rat lung tissues were washed and then homogenized on ice with normal saline. Homogenates from rat lung tissues or GFP-LC3/CCL149 cell culture supernatants were centrifuged at 12000rpm for 10min at 4 C, and the supernatants (100μL) were used for analysis. The levels of tumor necrosis factor-α (TNF-α), interleukin-1β (IL-1β), interleukin-10 (IL-6), macrophage chemoattractant protein-1 (MCP-1) and intercellular adhesion molecule-1 (ICAM-1) were measured by enzyme-linked immunosorbent assay (ELISA) kits (Elabscience, USA) in triplicate according to the manufacturer’s recommended protocol.

### Statistical Analysis

All data are presented as the mean± s.d. from at least three independent experiments. Student’s two-tailed *t*-test was used to compare the means of two-group samples. Two-way analysis of variance (ANOVA) was applied for the comparison of multiple groups in different H/R times. A one-way analysis of variance (ANOVA) was applied to determine the significant effect of 3-MA or rapamycin on studied rats pretreated with lung I/R followed by the least significant difference (equal variances assumed) or Tamhane’s T2 (equal variances not assumed) tests. All statistical analyses were performed using the *Graph Pad Prism5* software. *P* values less than 0.05 were considered significant. No statistical method was used to predetermine sample size. Randomization and a blinding strategy were used whenever possible.

## Competing interests

The authors declare no conflict of interest.

## Acknowledgements

We are grateful to the model organism collaborative innovation center of Wuhan University for its support with homologous male rats. We are grateful to the instruments of the school of life sciences at Wuhan University. This work was supported by award from the National Science Foundation of China Grants 81770095(to Q.G.), 81170076(to Q.G.), and 81700093(to T.F.).

